# Exploring Category Change Neural Responses Using Semantically Similar Stimuli With Fast Periodic Visual Stimulation

**DOI:** 10.64898/2026.06.22.733603

**Authors:** Zack Murphy, David Vandenheever

**Author notes:** Corresponding author contact information: Zack Murphy.

## Abstract

Fast periodic visual stimulation (FPVS) has recently demonstrated an ability to index multiple cognitive domains such as facial expression processing, working memory, and more recently, semantic categorization in just a few minutes of recording time. The present work investigates the effects of low semantic distance and how this modulates semantic categorization responses. Twenty-seven healthy young adults completed an FPVS oddball paradigm in order to determine whether comparing fruits and vegetables of the same color would elicit semantic categorization responses. Strong oddball responses were observed up to the 10^th^ frequency harmonic, and responses showed statistically significant right occipito-temporal lateralization that was not present in a low-level color change condition completed by participants and is opposite the pattern observed in recent word-based semantic categorization FPVS studies. The findings suggest that image-based FPVS paradigms should be investigated further as candidate tools to study conditions that affect semantic categorization such as Alzheimer’s disease.

## Introduction

Semantic categorization refers to one’s implicit ability to categorize items into groupings based on shared meaning, an ability that is especially important for interacting with our environment. For example, different dog breeds can vary widely in size, shape, but most people can easily classify them as dogs regardless of these variations. Similarly, dogs can easily be distinguished from a non-living object such as an apartment building with little effort. In people experiencing mild cognitive impairment, such as those diagnosed with Alzheimer’s disease (AD), their semantic categorization abilities can be significantly impaired (Stothart et al., 2021; David et al., 2025). Studying semantic categorization may therefore be useful in the detection and/or monitoring of conditions like AD that impair this process.

An increasingly popular methodology currently being used to study semantic categorization is fast periodic visual stimulation (FPVS) combined with electroencephalography (EEG). This methodology is based on the concept of steady state visually evoked potentials (SSVEPs), which describes that a visual stimulus presented at a fixed frequency (for example, 6 Hz) can elicit neural responses with a high signal-to-noise ratio (SNR) in the brain at that same frequency (Norcia et al., 2015). Because the frequency of the neural response is known prior to the start of the study, this response can be studied objectively and implicitly by recording neural activity and transforming it to the frequency domain using Fourier transforms. FPVS provides a quantifiable and objective signature of neural responses, and results that are sensitive on the individual level can be obtained in short recording times of just a few minutes per session (Retter & Rossion, 2016; Stothart et al., 2017; Volfart et al., 2021; David et al., 2025; Hermann et al., 2025).

Stimuli must be chosen carefully when designing an FPVS experiment, as the oddball response reflects the brain’s general discrimination between the base stimuli and oddball stimuli. As such, any visual difference between base and oddball images could influence the response. Hermann et al. (2025) demonstrated that the strength of the oddball response increased in correlation with the angle of rotation of a vertical line between base and oddball images, suggesting that oddball responses can be influenced by simple visual characteristics.

An unresolved issue in image-based semantic FPVS is whether category-change responses remain detectable when semantic distance and low-level visual differences are both reduced. Prior image-based FPVS studies have reported robust category-change responses for broad contrasts, but these contrasts often covary with shape, texture, color, and other low-level visual features. Word-based FPVS reduces image-level confounds and has produced reliable semantic categorization responses, including showing reduced responses in patients with AD (David et al., 2025), but word and image stimuli may not provide interchangeable cognitive access routes to semantic representations. Neuroimaging studies have shown modality-specific access regions for words and pictures, with word-semantic processing preferentially recruiting left temporal and inferior frontal language-related regions, and picture-semantic processing engaging posterior inferior temporal and lateral occipital regions more strongly (Vandenberghe et al., 1996). In addition, intracranial and cortical-stimulation studies indicate that reading and visually guided naming are partially dissociable processes within the ventral occipitotemporal cortex (Forseth et al., 2018; Woolnough and Tandon, 2024).

Additionally, semantic distance (how similar one object is in abstract category in relation to another object) may influence the way participants respond to stimuli; this was specifically recognized in the limitations of David et al. (2025), where the authors stated *“Finally, semantic distance between the two categories could also be modified, with the hypothesis that alternate words **more closely semantically related** to the repeated semantic category could be even more complicated to categorize for AD patients allowing to detect subtle semantic impairment at earlier disease stages.”*

Prior FPVS semantic categorization work with visual objects has typically relied on category contrasts that are broader than the present fruit versus vegetable design. Using picture stimuli, Stothart et al. (2017) reported reliable oddball responses for natural versus non-natural objects, animals versus non-animal natural items, and birds versus non-bird mammals. Milton et al. (2020) similarly showed that semantic category changes embedded periodically in a 6 Hz image stream elicited robust responses in both younger and older adults. More broadly, frequency-tagging studies have also shown reliable categorization of living versus non-living objects from infancy through adulthood, indicating that broad semantic distinctions are particularly tractable for FPVS paradigms (Peykarjou et al., 2024). Together, these findings demonstrate that semantic categorization can be measured robustly with FPVS, but they also indicate that some of the strongest prior effects have been obtained with category contrasts that differ substantially in both semantic class and overall visual statistics.

The oddball response in an image-based FPVS design reflects the brain’s discrimination of whatever properties systematically distinguish the base and oddball streams, not necessarily semantic category alone. Stothart et al. (2017) attempted to reduce such confounds by converting their images to grayscale and matching them on size, pixel intensity, and contrast, yet they still observed residual responses in scrambled control conditions and suggested that features such as feathers versus fur may not have been fully eliminated. Peykarjou et al. (2024) found that low-level image characteristics were sufficient to elicit reduced but measurable living/non-living categorization responses in phase-scrambled stimuli, while intact images generated markedly larger responses, likely because higher-order configurational information and category-level power-spectrum differences remained available. Other FPVS work makes the same point using simpler visual manipulations: oddball responses increase as line orientations deviate further from vertical (Hermann et al., 2025), contrast manipulations can strongly alter category-selective thresholds (Liu-Shuang et al., 2022), and viewpoint changes modify symmetry and feature visibility enough to produce view-specific neural effects in both FPVS paradigms (Or, Retter, & Rossion., 2021) and in older ERP based EEG work (Ewbank et al., 2008). These findings suggest that image-based semantic FPVS paradigms should be tightened as much as possible by limiting systematic low-level differences between base and oddball images. If a semantic categorization response remains under those conditions, the response can be interpreted with greater confidence as reflecting category-level processing rather than simpler visual deviance.

The first goal of the present work is to determine whether a semantic categorization response can be detected in healthy young adults even when the visual properties of the base and oddball stimuli are more tightly controlled. The second goal is to include a low-level visual oddball paradigm (see the discussion section of David et al., 2025) to act as a baseline against which we can compare semantic categorization responses. The present image-based FPVS study was designed within this context. To elicit semantic categorization responses, we displayed images of green fruits at 6 Hz, and images of green vegetables at 1.2 Hz, or every fifth image in the sequence, while limiting the low-level visual differences between images. The low-level visual condition included images of green vegetables as the base with red vegetables as the oddball. This was designed specifically to elicit a response to color changes without introducing a semantic category change. This approach was designed to limit the influence of shape, size, and color while simultaneously limiting the semantic distance, as fruits and vegetables are closely related but distinct categories. We hypothesize that even in this stimulus design in which visual properties between base and oddball images are more tightly controlled, a semantic categorization response will still be detectable in a sample of young healthy adults.

An additional strength of the present design is the inclusion of a low-level visual oddball condition as a direct comparison stream. David et al. (2025) argued that future FPVS studies in AD should test whether paradigms probing functions that are not typically impaired in AD, such as contrast or orientation oddballs, remain normal in the patient group. This recommendation is especially relevant because their word-based semantic categorization paradigm produced a strongly reduced semantic response in AD patients, reported as less than 25% of the response observed in healthy controls, while the use of words was intended specifically to minimize physical differences between stimulus categories. In this context, a low-level visual oddball stream provides a functional baseline against which semantic categorization responses can be interpreted and may help distinguish a selective semantic impairment from a more general deficit in generating oddball discrimination responses.

Such a control stream may also improve interpretation at the individual level. EEG amplitudes vary substantially across participants for reasons that are not purely cognitive, including skull and scalp properties and other volume-conduction factors; even subject position can change occipital EEG magnitude, and normalization procedures have been shown to improve evoked-potential reproducibility (Cuffin, 1993; Rice et al., 2013; You et al., 2012). For that reason, an exploratory ratio relating the summed baseline-corrected amplitude (BCA) from the low-level oddball condition to the semantic-categorization BCA could provide a useful within-subject normalization index. In the case of AD-like impairment, a participant might show a relatively preserved low-level oddball response but a diminished semantic categorization response, yielding an abnormally elevated low-level/semantic ratio.

## Materials and Methods

### 2.1 Participants

Twenty-seven participants were recruited from the Mississippi State University student population with a mean age of 20.3 years, SD 1.22, and 8 of the 27 were male. No a priori power analysis was conducted. The target sample size was therefore determined prior to the beginning of the study based on available time, resources, and sample sizes used in previously published FPVS work: n = 12 in Rossion et al., 2015, n = 14 in Volfart et al., 2021, n = 16 in Retter & Rossion, 2016, and n = 20 in Stothart et al., 2017. The final sample of 27 participants in the present study was therefore consistent with, and larger than, many sample sizes used in recently published FPVS work. All participants reported normal or corrected to normal vision and no color blindness. Additionally, they were provided written informed consent prior to the beginning of the study and were free to withdraw at any time. Participants were not compensated for the study. This study was approved by the Mississippi State University Institutional Review Board.

### 2.2 Stimuli

Stimuli included photorealistic images of culinary fruits and vegetables presented on plain gray backgrounds that varied in color depending on experimental condition. All images were 256 x 256 resolution and were displayed at a visual angle of 5 degrees in height by 5 degrees in width corresponding with a viewing distance of 80 cm from a 24-inch screen with a resolution of 1920 x 1080. Each image in the stimulus set was generated using Midjourney, an AI-based image generation tool. Each image was generated using a variation of the following prompt: “Generate a photorealistic image of a [(color) fruit/vegetable] on a plain gray background with nothing else in the image. All images should be 256 x 256 resolution, and the camera angle should be a frontal, head on angle.”

Following image generation, images were manually inspected and selected for inclusion in the study. Images with obvious image generation artifacts, images that varied in shape or size from the intended goal, and images that did not appear photorealistic were excluded from the final image set. Images were also selected to reduce large differences in object size and position within the 256 x 256 frame across the set. For the color-change condition (see Figure 1), images were selected to keep color relatively consistent within each color group (for example, each image of a red vegetable was selected so that the red coloration was visually similar to that of the other red vegetable in the stimulus set). For the semantic categorization condition (see Figure 2), the images were selected to maximize similarity between the green fruit and green vegetable image sets in color, orientation, and general shape. The image dataset included 250 total images (125 images, with 100 base images and 25 oddball images per condition). In the semantic response condition, the green fruit base set included ten different fruit categories with ten images in each category: green apples, limes, avocados, green bananas, green grapes, honeydews, green pears, watermelon, kiwis, and green plums. The green vegetable oddball set included five categories: broccoli, cabbage, green bell peppers, jalapenos, and spinach, with five exemplars per category.

**Figure 1.**
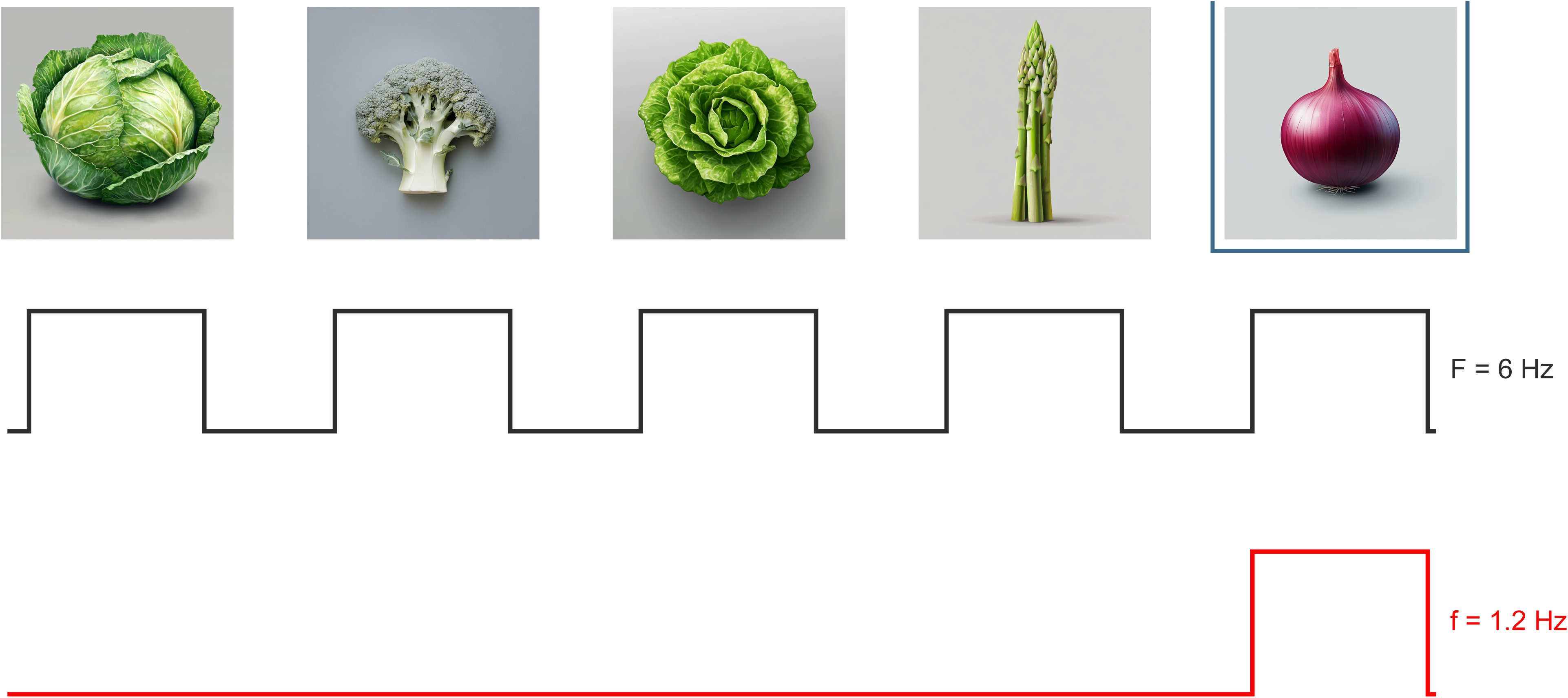
illustrates one cycle of the FPVS oddball condition designed to elicit a color response. The first four images in the sequence are green vegetables, and the fifth image in the sequence is a red vegetable. This cycle repeats at 6 Hz for 147 total cycles.

**Figure 2.**
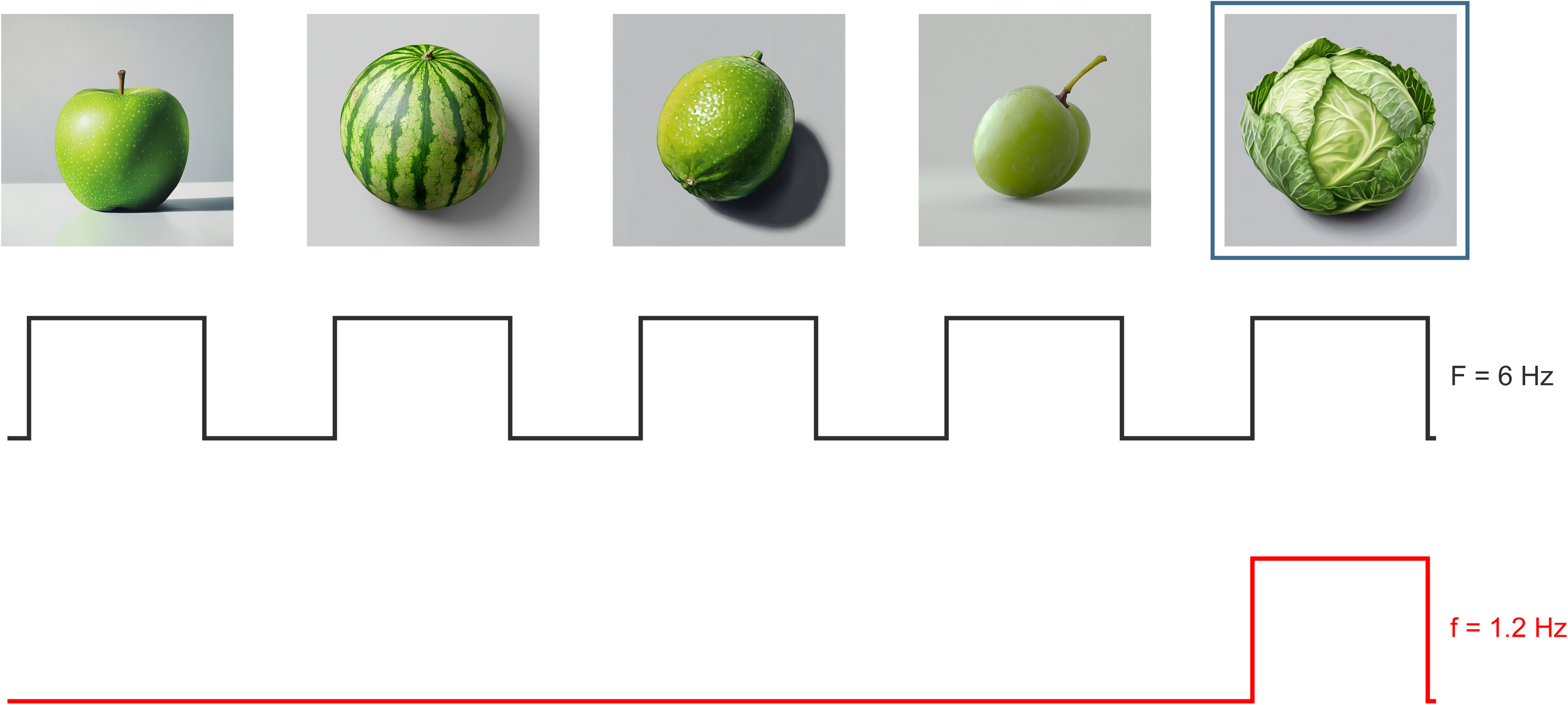
illustrates one cycle of the FPVS condition designed to elicit a semantic categorization response. The first four images in the sequence are green fruits while the fifth image of every sequence is a green vegetable.

In the color response condition, green vegetables served as base images and red vegetables served as oddballs. The green vegetable base set included broccoli, green bell peppers, jalapenos, asparagus, green beans, cabbage, green peas, bok choy, leafy greens, zucchini, and spinach. The red-vegetable oddball set included red bell peppers, radishes, red onions, red chili peppers, and tomatoes. Across 147 five-image cycles, each cycle contained four base images and one oddball image, yielding 588 base-image presentations and 147 oddball presentations per condition. Each base image and each oddball image was presented approximately six times per condition with no immediate repeats.

### 2.3 Procedure

#### 2.3.1 Experimental Design

This experiment used an FPVS oddball design with two experimental conditions. In each condition, base images were presented at 6 Hz, and oddball images were presented at 1.2 Hz (every fifth image in the stream). Images were displayed on screen for 83 ms followed by an 83 ms blank inter-stimulus interval before the next image was shown. Conditions included 147 cycles of the five-image stream. Both conditions lasted roughly 122 seconds and were presented to the participant in random order. The primary condition of interest was the condition designed to elicit a semantic categorization response. To do this, images of green fruits were used as base stimuli and images of green vegetables were used as oddball stimuli. Stimuli were chosen to lower both the semantic distance and the visual variability between base and oddball images. The color response condition included images of green vegetables as base images and red vegetables as oddball images. To maintain participant attention, they were instructed to focus their gaze on a blue fixation cross at the center of the screen (Rossion et al., 2015). The fixation cross changed colors from blue to red at random times during each condition for a total of seven color changes in each condition. Participants were instructed to press the space bar whenever they detected this change to ensure their attention remained constant throughout the study.

#### 2.3.2 EEG Recording

Brain activity was recorded using a BioSemi ActiveTwo EEG system (BioSemi, Amsterdam, The Netherlands) using a 64-electrode cap arranged according to the international 10-10 system. Two external electrodes were placed on the left and right mastoids as references. All data were sampled at 2048 Hz.

### 2.4 EEG Analysis

#### 2.4.1 Data processing

All data were processed using a custom MNE Python-based FPVS data analysis pipeline using Python 3.13.5 following established FPVS data processing techniques (see Rossion et al., 2020, Volfart et al., 2021, Hermann et al., 2025, David et al., 2025). Raw data were imported individually for each participant and initially referenced using two external mastoid electrodes. A basic finite impulse response (FIR) band-pass filter from 0.1 Hz to 50 Hz was applied. Data were then downsampled from 2048 Hz to 256 Hz to reduce file size and processing time. Noisy EEG channels were then automatically identified using channel-level kurtosis. Kurtosis values were calculated for each channel and converted to z-scores relative to the distribution of channels in that recording. To reduce the influence of extreme values on the channel distribution, the highest and lowest 10% of kurtosis values were removed before calculating the mean and standard deviation used for z-score conversion. Channels that exceeded 5 standard deviations from this mean were interpolated using spherical spline interpolation. Across the final dataset, an average of 2.78 electrodes were interpolated per participant, corresponding to an average of 4.27% of the 64 electrode montage. After interpolation, data were re-referenced to the common average reference. As in Vandenheever et al. (2025), artifact rejection methods (e.g., Independent Component Analysis) were not applied.

The data were then segmented by extracting each experimental condition into separate files. Each condition was further cropped to ensure that each segment contained the same integer number of oddball cycles (147 cycles). Following preprocessing, each EEG segment was transformed into the frequency domain using a Fast Fourier Transform (FFT).

#### 2.4.2 Defining Significant Harmonics

For each target frequency bin, baseline-corrected amplitude (BCA), signal-to-noise ratio (SNR), and z-scores were calculated from the amplitude spectrum. BCA was calculated by subtracting the mean amplitude of the surrounding local-noise bins from the raw amplitude at the target frequency. SNR was calculated by dividing the raw amplitude at the target frequency by the same local-noise estimate. The local-noise window included 10 frequency bins below and 10 frequency bins above the target frequency, excluding the target bin and the immediately adjacent bins. The minimum and maximum finite values within the local-noise window were removed before calculating the local-noise mean. Z-scores were calculated as the difference between the target amplitude and the local-noise mean, divided by the standard deviation of the local-noise bins. Harmonics overlapping with the 6-Hz base stimulation frequency or its harmonics were excluded. Harmonic selection was performed once at the group level using the grand-averaged amplitude spectrum across included participants, analyzed conditions, and scalp electrodes, after excluding the base rate frequency of 6 Hz and its harmonics. A z-score was calculated at each candidate oddball harmonic, and harmonics were considered significant when z > 1.64, corresponding to a one-tailed p < .05 threshold. The same list of significant harmonics was applied to every participant, condition, and ROI. BCA values were summed across significant oddball harmonics and averaged across electrodes within each ROI (Retter et al., 2021). The resulting ROI level BCA values were used for statistical analysis.

#### 2.4.3 Regions of Interest

Three regions of interest (ROIs) were defined a priori based on the scalp distributions reported in prior FPVS studies of semantic categorization and adapted to our 64 electrode BioSemi 10-10 montage. The left occipito-temporal (LOT) ROI included P7, P9, PO7, PO3, and O1, and the right occipito-temporal (ROT) ROI included P8, P10, PO8, PO4, and O2. These ROIs were included because semantic FPVS responses have been reported over left occipito-temporal electrodes. We included the right occipito-temporal ROI for comparison (Volfart et al., 2021; David et al., 2025). A central ROI (FCz, Cz, CPz, CP1, C1, FC1) was also included because David et al. (2025) reported a semantic categorization response over this ROI.

### 2.5 Statistical Analysis

Statistical analysis was performed on summed ROI-level BCA values in Python 3.13.5 using custom Python scripts. The primary ROI-level analysis tested whether summed BCA values exceeded zero in the Semantic Response and Color Response conditions. Response-vs-zero tests were performed separately for the LOT, ROT, and Central ROIs. Normality was evaluated using Shapiro-Wilk tests. When normality assumptions were met, two-tailed one-sample t-tests were used; when they were not met, two-tailed Wilcoxon signed-rank tests were used. P-values for planned response-vs-zero tests were Holm-corrected (Holm, 1979) across ROIs within each condition. All planned ROI tests used α = .05 and were evaluated using adjusted p-values. Cohen’s dz was reported for t-tests. Planned Semantic Response versus Color Response contrasts were tested within each ROI and Holm-corrected.

Lateralization between left and right hemispheres was evaluated by comparing mean summed BCA values in LOT and ROT ROIs. Values were computed as ROT – LOT. Positive values indicated that the response was stronger in ROT, while negative values indicated the opposite. This contrast was applied to both the semantic response and the color response conditions. Planned lateralization contrasts were Holm-corrected.

### 2.6 Source estimation

L2 minimum-norm source estimation was performed at the individual level on the EEG data using MNE python. The inverse solution used no noise normalization, no depth weighting, and a loose orientation constraint of 0.2, and all other implementation methods were followed as suggested in Hauk et al. (2021) and Hauk et al. (2025). For each participant and condition, source estimates were baseline-corrected and converted to z-scores using the same procedure described in 2.4.2. We used the FreeSurfer fsaverage 6 MRI template (https://nilearn.github.io/dev/modules/description/fsaverage.html) with an ico3 cortical surface for all participant source localization mapping as no individual MRI templates were available. The resulting group average data was painted onto the fsaverage templates for visualization (see Figure 7).

## 3. Results

The semantic response was significantly greater than zero in all ROIs after Holm correction, confirming this study’s hypothesis that semantic responses can be detected even when semantic distance is low and visual variability is limited. In Left OT, mean BCA was 0.44 µV, SD = 0.27, t(26) = 8.50, p < .001, d_z_ = 1.64. In Right OT, mean BCA was 0.667 µV, SD = 0.333, t(26) = 10.40, p < .001, d_z_ = 2.00. In the central ROI, mean BCA was 0.24 µV, SD = 0.18, t(26) = 6.77, p < .001, d_z_ = 1.30.

The largest semantic response amplitude was observed in Right OT, and this semantic response was significantly larger in Right OT than Left OT (mean Right OT–Left OT difference = 0.228 µV, t(26) = 4.04, p = .001, d_z_ = 0.78; see Figures 3 and 4). Notably, this ROT lateralization was not significant in the color condition (mean BCA difference between ROT and LOT was 0.034 µV, Wilcoxon *p* = .470, not significant). Grand average scalp topography plots (see Figure 6) and source estimation analysis (see Figure 7) also show ROT lateralization.

**Figure 3:**
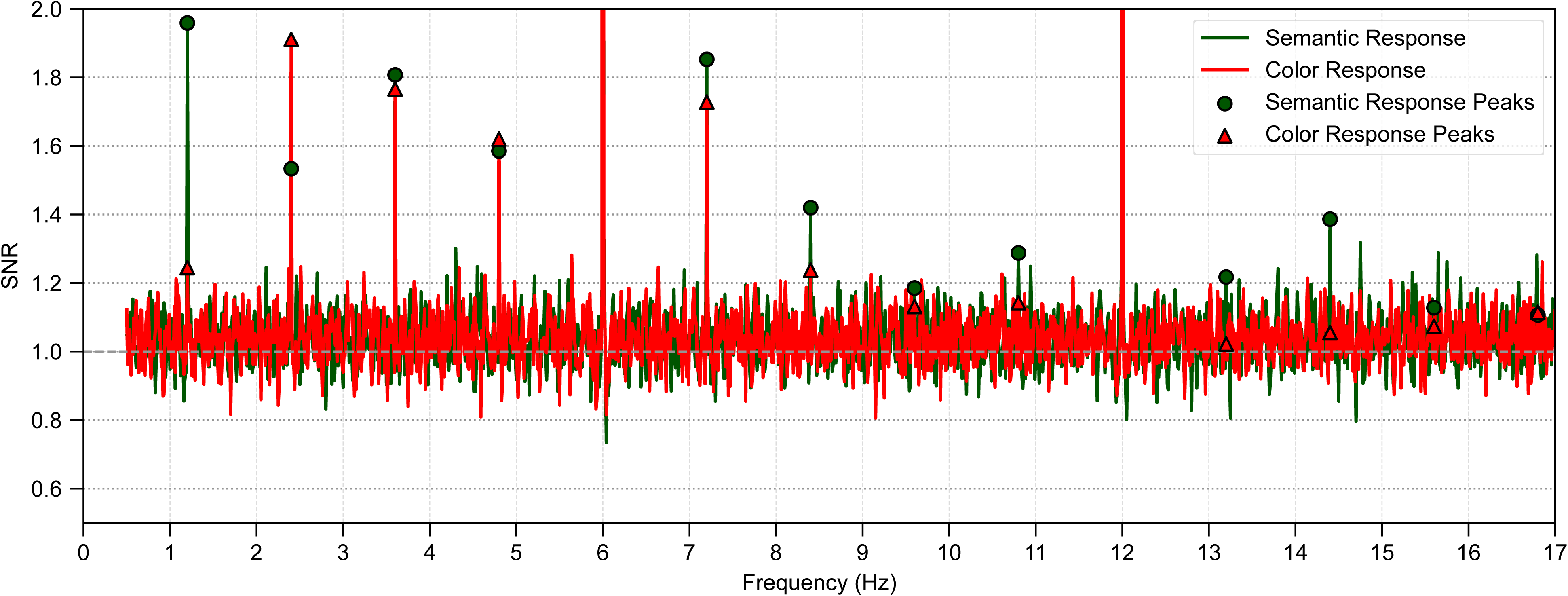
Signal-to-noise ratio (SNR) plot of the responses in the right occipito-temporal region. Oddball responses to the semantic condition are plotted in green, and responses to the color change condition are plotted in red. Note that an SNR value of near 2.0 indicates a near 100% increase in the signal strength relative to background noise.

**Figure 4:**
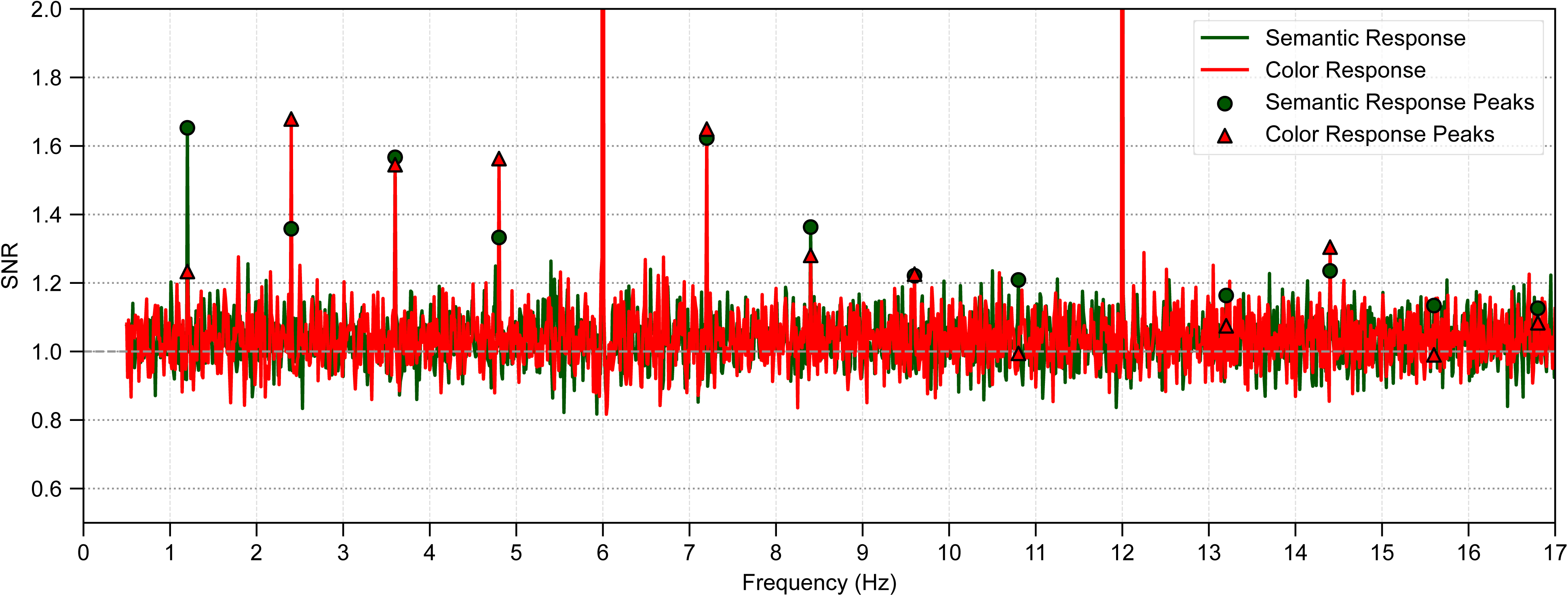
Signal-to-noise ratio (SNR) plot of the responses in the left occipito-temporal region. Oddball responses to the semantic condition are plotted in green, and responses to the color change condition are plotted in red.

The low-level Color Response condition also elicited significant summed BCA responses in all reported ROIs after Holm correction. In Left OT, mean BCA was 0.41 µV, SD = 0.33, t(26) = 6.41, p < .001, d_z_ = 1.23. In Right OT, Mean BCA was 0.442 µV, SD = 0.524. Because the Right OT Color Response failed the Shapiro-Wilk normality test, the Wilcoxon signed-rank test was selected and the response remained significant after Holm correction, p < .001. In Central, Mean BCA was 0.16 µV, SD = 0.18, t(26) = 4.55, p < .001, d_z_ = 0.88.

The Right OT - Left OT contrast in the color response was small and non-significant (mean Right OT–Left OT difference = 0.034 µV, selected Wilcoxon p = .470).

SNR plots show strong neural responses to both conditions in ROT (see Figure 3), LOT (see Figure 4) and the Central ROI (See Figure 5).

**Figure 5:**
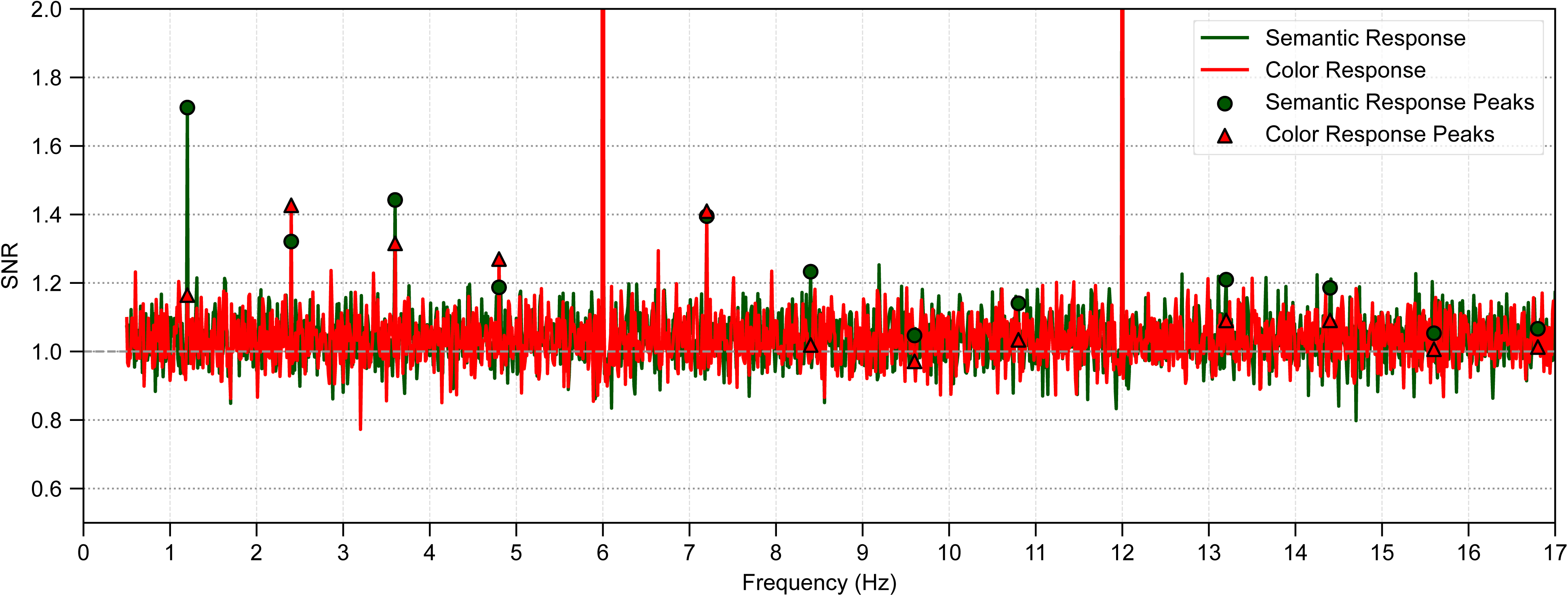
Signal-to-noise ratio (SNR) plot of the responses in the central region. Oddball responses to the semantic condition are plotted in green, and responses to the color change condition are plotted in red.

**Figure 6:**
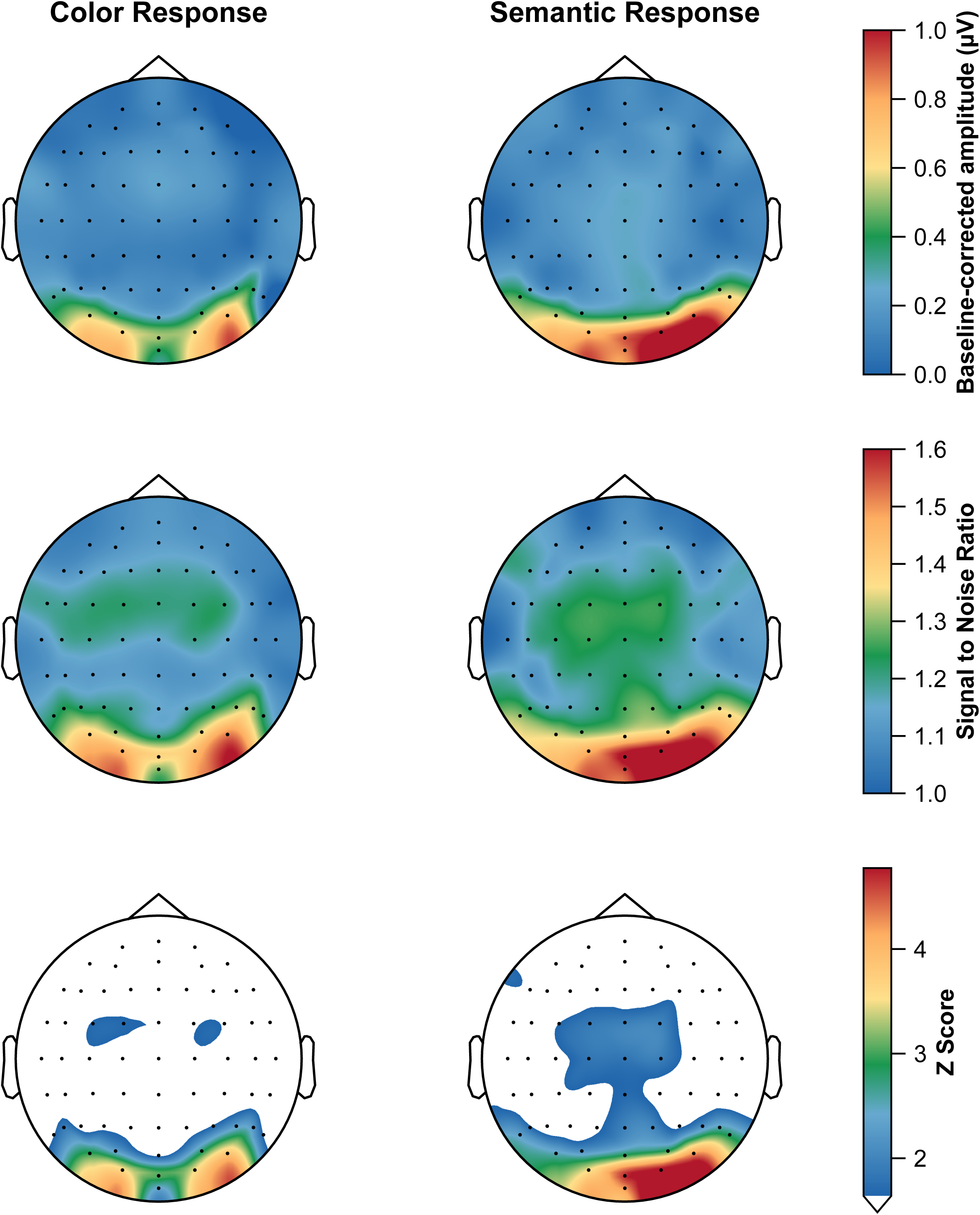
Top-down illustrations of the scalp averaged data across all participants in the color response and semantic response conditions. Note the semantic categorization condition is more prominent over the right occipito-temporal ROI. White values correspond to non-significant values with a z-score below z = 1.98.

### 3.1 Source Estimation Results

Source space z-score maps are shown in Figure 7. The figure was masked with a cluster-permutation correction to ensure only responses significant across the group were plotted (Hauk et al., 2021; Hauk et al., 2025). Source estimation visualization independently revealed the same ROT lateralization pattern that was observed via plotting and analyzing baseline corrected amplitude values.

**Figure 7:**
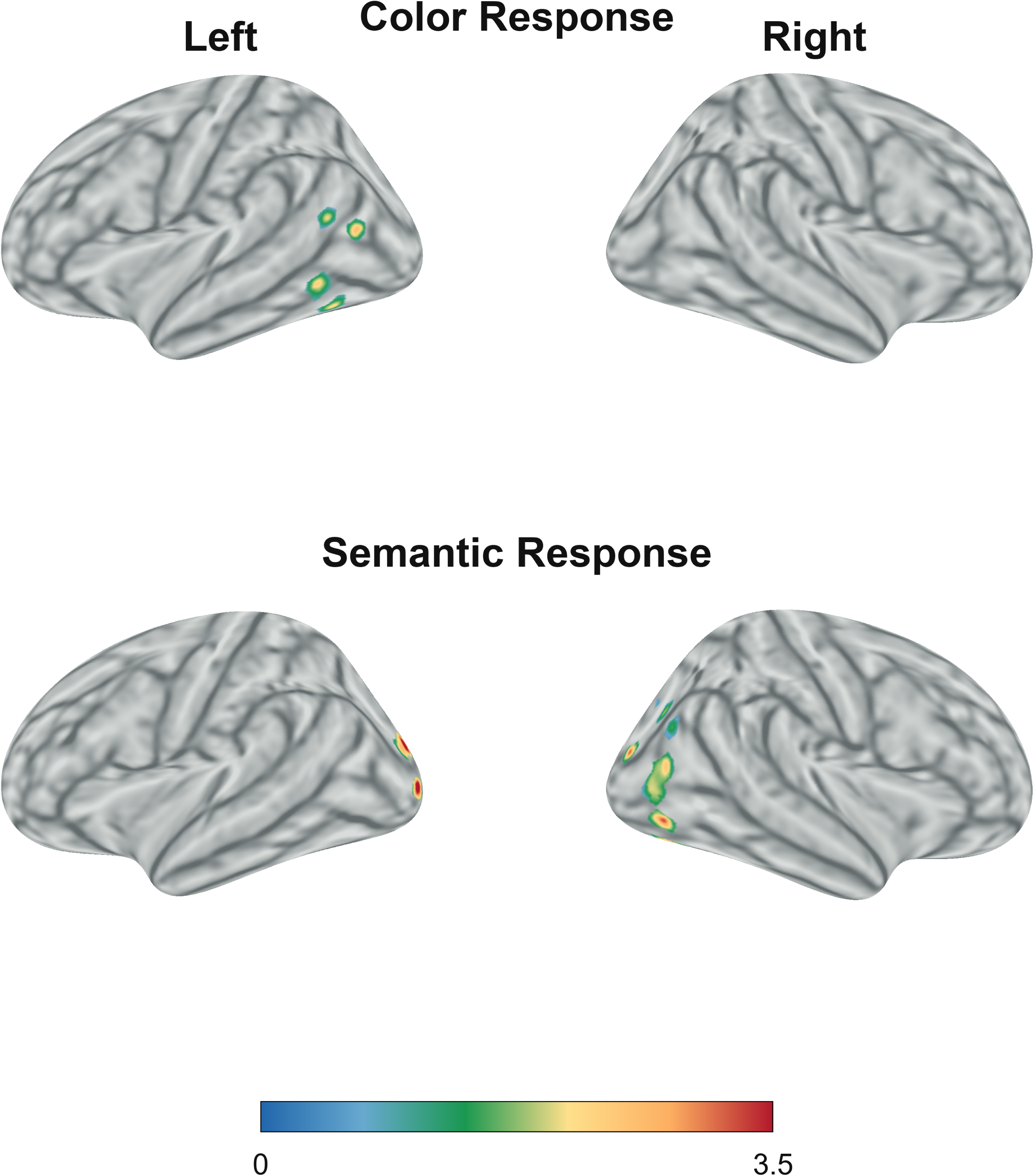
L2 source estimation plots of both the summed z-score values in the color and semantic response conditions on the left and right hemisphere. A cluster-based permutation mask was applied; non-significant vertices at the group level are masked and displayed in gray. Note that the group average response to the color condition in the right hemisphere shows no significant response, while the semantic response is primarily present in the right hemisphere.

### 3.2 Individual level responses

Participants were examined at the individual level as in David et al. (2025). Detectability at the individual level was examined at both the ROI level and at the individual electrode level. At the ROI level, the semantic response was significant (exceeding *Z* > 1.64, *p* < .05) in 23 of 27 participants over Right OT, 21 of 27 over Left OT, and 18 of 27 over Central. At a threshold of *Z* > 2.32 (*p* < .01), the semantic response remained detectable in 22 of 27 participants over Right OT, 18 of 27 over Left OT, and 12 of 27 over the central ROI. At *Z* > 3.10 (*p* < .001), 19 of 27 participants retained a significant response over Right OT, 14 of 27 over Left OT, and 7 of 27 over central.

For the color response condition, ROI-level detectability at *Z* > 1.64 was observed in 18 of 27 participants in each ROI. At *Z* > 2.32, the response was detected in 15 of 27 participants over Right OT, 14 of 27 over Left OT, and 11 of 27 over Central. At *Z* > 3.10, 14 of 27 participants retained a significant response over Right OT, 12 of 27 over Left OT, and 3 of 27 over Central.

We also quantified individually significant responses using electrode-level significance across electrodes within each participant following BH-FDR correction (Benjamini & Hochberg, 1995) following electrode level detectability thresholds described in David et al. (2025). For the semantic response condition, 26 of 27 participants had at least one FDR-significant electrode across the whole scalp, with a mean of 21.44 ± 15.59 significant electrodes per participant. FDR-significant electrodes were observed in 23 of 27 participants over Right OT, 21 of 27 over Left OT, and 17 of 27 over Central. For the Color Response condition, 22 of 27 participants had at least one significant electrode across the whole scalp, with a mean of 16.63 ± 15.09 significant electrodes per participant. At the ROI level, significant electrodes were observed in 20 of 27 participants over Right OT, 20 of 27 over Left OT, and 17 of 27 over Central.

### 3.3 Base rate response

General visual responses were observed at the 6-Hz base rate and harmonics in the bilateral OT ROI. For the Semantic Response condition, the base-rate response exceeded *Z* > 1.64 through 48 Hz: 6 Hz, *Z* = 25.88, *p* < .001; 12 Hz, *Z* = 9.15, *p* < .001; 18 Hz, *Z* = 9.63, *p* < .001; 24 Hz, *Z* = 6.57, *p* < .001; 30 Hz, *Z* = 3.69, *p* < .001; 36 Hz, *Z* = 2.41, *p* = .008; 42 Hz, *Z* = 1.82, *p* = .034; and 48 Hz, *Z* = 1.80, *p* = .036. For the Color Response condition, the base-rate response exceeded *Z* > 1.64 through 36 Hz: 6 Hz, *Z* = 26.79, *p* < .001; 12 Hz, *Z* = 9.11, *p* < .001; 18 Hz, *Z* = 9.11, *p* < .001; 24 Hz, *Z* = 4.82, *p* < .001; 30 Hz, *Z* = 2.46, *p* = .007; and 36 Hz, *Z* = 2.12, *p* = .017.

### 3.4 Exploratory ratio metric

As an exploratory within-participant normalization metric, the BCA from the semantic response was divided by the color response BCA values within each ROI. Ratios near 1.0 indicate that the semantic response and color response were similar in amplitude at the group level. Median semantic/color ratios were close to 1.0 in all reported ROIs. In Central, the raw ratio had a median of 1.00, mean = 1.20, SD = 3.43, range = -2.50 to 16.81. After trimming the single minimum and maximum ratio, the raw central ratio remained centered near 1.0, median = 1.00, mean = 0.73, SD = 1.33, range = -2.34 to 3.32. In Left OT, the raw ratio had a median of 0.98, mean = 1.06, SD = 1.16, range = -2.33 to 3.06. After trimming, the Left OT ratio had a median of 0.98, mean = 1.12, SD = 0.90, range = -0.67 to 2.94. In Right OT, the raw ratio had a median of 1.13, mean = 0.93, SD = 7.72, range = -25.81 to 26.24. After trimming, the Right OT ratio remained close to 1.0 by central tendency, median = 1.13, mean = 0.99, but variability remained high, SD = 2.83, range = -11.19 to 5.59.

Across ROIs, the median ratios suggest that the Semantic Response was approximately comparable in magnitude to the Color Response in this young-adult sample. However, participant-level ratios were variable. The proportion of participants within ±20% of the ROI median was 30% in Central, 19% in Left OT, and 30% in Right OT.

## Discussion

The present study is, to our knowledge, the first to demonstrate that semantic categorization responses can be detected using visual FPVS paradigms under low semantic distance and low visual variability constraints. In contrast to previous literature (e.g., the comparison of natural objects versus man mange objects, or animals versus cities) the present fruit versus vegetable constraint provides a stricter test of visual semantic categorization than has previously been explored.

A second key result was that semantic responses were largest over right occipito-temporal electrodes and significantly greater in the ROT region versus the LOT. This pattern of lateralization was not observed in the low-level color change condition, providing evidence that the observed pattern is specific to semantic categorization and is not a product of general rapid image presentation. Additionally, these results suggest that image based semantic categorization may follow a different neural pattern than word based semantic categorization, as previous word-based FPVS research shows that word-based paradigms elicit stronger responses over LOT (David et al., 2025; Hauk et al., 2025; Volfart et al., 2021). These studies established that FPVS can measure semantic categorization in short recording times, but they did not examine the effect of semantic distance on categorization responses. One plausible explanation for the observed pattern is that semantic categorization measured with visual FPVS based paradigms reflects partly different mental access routes than previous word based semantic categorization paradigms. This interpretation is consistent with prior work suggesting that word versus picture based semantic access relies on similar but separable cognitive pathways, including stronger engagement of left lateralized language recruitment for words (Vandenberghe et al., 1996; Forseth et al., 2018; Woolnough & Tandon, 2024). The observed semantic response may reflect a visual dominant conceptual processing route that is expressed most strongly over right occipito-temporal regions (similar to the face selective responses found in earlier face based FPVS work, e.g., Rossion et al., 2015) whereas word-based paradigms tend to elicit responses in left-lateralized language systems.

The potential clinical utility of semantic categorization FPVS has been discussed in previous publications (e.g., see Stothart et al., 2017, 2020, and 2025) and we recommend the same. FPVS has several key advantages over traditional ERP based methods (short recording times, high SNR, individual level detectability, smaller group sizes), over other neuroimaging methods (short recording times, low cost) and over behavioral tasks (objective neural measurements, no participant task instructions required) for studying cognitive decline, and with further research into semantic based paradigms, FPVS could prove to be quite useful in the early detection of Alzheimer’s disease and aid in earlier diagnoses. However, despite the strengths of this design and the promise of future utility, several limitations of the present work should be acknowledged.

First, the design of the stimuli in this condition was done with Midjourney, an AI based image generation tool. The motivation of using this tool to generate images was twofold: first, we wanted to validate whether AI generated images were feasible for use in FPVS paradigms, and second, we wanted more visual control over the images. Using an image generation tool allowed us, within reason, to control the visual properties of each image in the stimulus set and to build an acceptable image database of images quickly. However, the use of this image generation method did result in slightly different colors, background contrasts, and sizes of the stimuli, so future studies should attempt to further control contrast, luminance, and relative size of the fruits and vegetables within the stimuli.

Second, the premise of this study assumes that the participant has a strong conceptual understanding of the difference between fruits and vegetables in order to detect a semantic category change from fruit to vegetable. For example, certain foods like tomatoes are often considered a culinary vegetable, even though they are technically a fruit, so tomatoes were excluded from the stimulus set. However, it is still possible that some participants may have had a varying concept of what foods fall into the culinary fruit or the vegetable category, which could have affected the results. Future studies with similar designs may incorporate priming or norming to encode which foods are fruits and which foods are vegetables prior to the beginning of the study, or use other stimulus sets with similar semantic distance that may be more universally understood.

Third, while our green fruit versus green vegetable condition was designed specifically as a low semantic distance condition, we did not introduce a similar visually constrained condition with high semantic distance. Future studies should consider including a high semantic distance condition to explore the effects of how modulating this parameter affects FPVS responses.

Finally, the spatial resolution of EEG is a limitation of any study of this nature. While we did utilize L2 source estimation methods (Hauk et al., 2021; Hauk et al., 2022; Hauk et al., 2025) which provided a separate and independent measure of neural activity on the cortical surface, this method does not address the issue of source localization deeper within the brain, and cannot directly solve the spatial resolution problem of EEG. Future studies should consider combining EEG with higher resolution methods such as MEG or fMRI and employing source localization methods such as LORETA to attempt to account for this.

## Conclusion

We provide evidence that semantic categorization can be measured in healthy young adults using a sensitive FPVS design with limited visual variability and low semantic distance. This approach may be useful for future research in clinical settings with patients who are experiencing AD-like symptoms.

## Acknowledgements

During the preparation of this work, the authors used Codex, a desktop-based coding tool from OpenAI that uses ChatGPT Pro 5.5 for debugging custom scripts used in data analysis and figure generation in this manuscript. Additionally, images in the stimulus set were generated using Midjourney, an image generation tool, as part of the key design principles of this study The authors manually reviewed and edited the code, figures, and images used in this research as needed and take full responsibility for the content of the published article. The authors received no funding of any kind from any organization to complete this work and declare no competing interests.

